# A Novel *Ex Vivo* Approach for Investigating Profibrotic Macrophage Polarization Using Murine Precision-Cut Lung Slices

**DOI:** 10.1101/2024.07.05.602278

**Authors:** Megan Vierhout, Anmar Ayoub, Pareesa Ali, Vaishnavi Kumaran, Safaa Naiel, Takuma Isshiki, Joshua F. Koenig, Martin R.J. Kolb, Kjetil Ask

## Abstract

Idiopathic pulmonary fibrosis (IPF) is fatal interstitial lung disease characterized by excessive scarring of the lung tissue and declining respiratory function. Given its short prognosis and limited treatment options, novel strategies to investigate emerging experimental treatments are urgently needed. Macrophages, as the most abundant immune cell in the lung, have key implications in wound healing and lung fibrosis. However, they are highly plastic and adaptive to their surrounding microenvironment, and thus to maximize translation of research to lung disease, there is a need to study macrophages in multifaceted, complex systems that are representative of the lung. Precision-cut lung slices (PCLS) are living tissue preparations derived from the lung that are cultured *ex vivo,* which bypass the need for artificial recapitulation of the lung milieu and architecture. Our objective was to establish and validate a moderate-throughput, biologically- translational, viable model to study profibrotic polarization of macrophages in the lung using murine PCLS. To achieve this, we used a polarization cocktail (PC), consisting of IL-4, IL-13, and IL-6, over a 72-hour time course. We first demonstrated no adverse effects of the PC on PCLS viability and architecture. Next, we showed that multiple markers of macrophage profibrotic polarization, including Arginase-1, CD206, YM1, and CCL17 were induced in PCLS following PC treatment. Through tissue microarray-based histological assessments, we directly visualized and quantified Arginase-1 and CD206 staining in PCLS in a moderate-throughput manner. We further delineated phenotype of polarized macrophages, and using high-plex immunolabelling with the Iterative Bleaching Extends Multiplexity (IBEX) method, showed that the PC effects both interstitial and alveolar macrophages. Substantiating the profibrotic properties of the system, we also showed expression of extracellular matrix components and fibrotic markers in stimulated PCLS. Finally, we demonstrated that clodronate treatment diminishes the PC effects on profibrotic macrophage readouts. Overall, our findings support a suitable complex model for studying ex vivo profibrotic macrophage programming in the lung, with future capacity for investigating experimental therapeutic candidates and disease mechanisms in pulmonary fibrosis.

## INTRODUCTION

Fibrotic interstitial lung diseases (ILD) are a group of debilitating disorders characterized by excessive scarring of the lung tissue and declining respiratory function. Idiopathic pulmonary fibrosis (IPF) is one of the most common ILD subtypes, for which the cause remains unclear and prognosis is relatively short, with patients surviving a median of 3 to 5 years after diagnosis [1]. There are only two approved antifibrotic therapies for IPF, nintedanib and pirfenidone. Despite slowing the progression of disease, neither are curative or reverse fibrosis [2,3]. Given the poor prognosis and limited treatment options for fibrotic lung disease, investigation is urgently needed to delineate pathogenesis, as well as to develop novel approaches to rapidly screen emerging experimental treatments and their respective effects on the lung microenvironment.

Macrophages, the most abundant immune cells in the lung, have vital implications in mediating the pathology of lung fibrosis [4,5]. Specifically, there is evidence supporting the contribution of profibrotic alternatively activated macrophages to aberrant wound healing [6–9], rendering these cells and their programming critical targets of interest in the context of lung fibrosis. Notably, macrophages are incredibly plastic cells with a dynamic phenotypic spectrum. They are highly interactive and swiftly adapt to their surrounding microenvironment, participating in a complex crosstalk of multifaceted signalling [9]. As such, evaluation of macrophages and their programming *in vitro*, although favoured for throughput and speed in preclinical screening studies, is limited for modelling authentic macrophage behaviour in the lung and fibrotic milieu. Therefore, to maximize translation to disease, there is a need to study macrophages in an environment that better represents the lung.

Precision cut lung slices (PCLS) are living tissue preparations derived from the lung that are cultured *ex vivo*. Historically, PCLS were used for pulmonary airway studies, mainly focusing on smooth muscle and airway epithelial cells [10,11]. More recently, research involving PCLS has become increasingly popular for the study of numerous cellular mechanisms, fibrosis, infection, inflammation, senescence, disease-relevant stimuli, and experimental treatments [12]. A primary advantage of the PCLS system is that slices contain all resident cells of the lung and maintain intercellular interactions and cell-to-matrix relationships which are not found in traditional two-dimensional cell culture or three-dimensional organoid systems [13]. Conventional systems face limitations related to physiologically accurate lung cell and extracellular matrix (ECM) localization patterns, which are overcome in PCLS as lung structures do not need to be artificially recapitulated [14]. The nature of the PCLS model is also less time- and resource-intensive than *in vivo* disease models, and in honouring the 3 Rs of Replacement, Reduction, and Refinement, markedly decreases the number of animals needed for experiments [15]. As numerous slices can be obtained from lungs of various mammals, PCLS have been leveraged for their high-throughput potential, especially for screening compounds [16–19]. Thus, there is a need to develop readouts that are also throughput-conducive to best maximize the potential of this platform.

Recent single-cell RNA sequencing (scRNAseq) studies have demonstrated the preservation of immune cells in PCLS [20], including proliferating macrophages, alveolar macrophages, and monocyte-derived macrophages. Despite being disconnected from the influence of the circulating immune system and thus incoming monocyte infiltration, Blomberg et al. (2023) have recently exhibited proliferative capacity of macrophages in their murine PCLS model of pre-cancer malignancy [21]. Additionally, Alsafadi et al. (2017) [22] successfully induced multiple fibrotic changes in human PCLS using a fibrotic cocktail (FC). Using this FC, Lang et al. (2023) recently showed increased expression of fibrosis-related marker genes in a population of SPP1^+^ macrophages in human PCLS [23]. Overall, this evidence offers the suggestion that PCLS may constitute a suitable medium for the study of macrophage programming in the lung, and further investigation is needed to study profibrotic polarization and modes to measure this.

Our group has previously demonstrated that the addition of IL-6 to the traditional alternative programming cocktail of IL-4 and IL-13 effectuated a hyperpolarized profibrotic phenotype in murine and human macrophages *in vitro* [24]. Therefore, we postulated that the profibrotic effects of this cocktail could be applied in an appropriate *ex vivo* system, potentially contributing to the development of a novel translational model for studying pulmonary profibrotic macrophages. Here, we evaluate the potential of employing this cocktail to establish and validate a moderate-throughput, complex, viable model to study profibrotic polarization of macrophages in the lung using PCLS. We demonstrate that treatment with IL-4+IL-13+IL-6, referred to as the polarization cocktail (PC), effectively induces profibrotic programming of *ex vivo* macrophages in murine PCLS over a 72-hour time course study. We implement the use of multiple readouts, with a focus on establishing throughput of the model, especially through quantitative histological readouts using tissue microarray-based approaches. Substantiating the profibrotic properties of the system, we also show expression of ECM and fibrotic markers in PCLS following PC treatment. Finally, we demonstrate that clodronate treatment diminishes the effects of the PC on profibrotic macrophage readouts.

## METHODS

### Animal Utilization

All work involving animals was approved by the McMaster University Animal Research Ethics Board (Animal Utilization Protocol #23-19) and was conducted in accordance with the Canadian Council on Animal Care guidelines. Wildtype female C57BL6/J mice (The Jackson Laboratory) aged 8–12 weeks were housed in pathogen-free conditions at the McMaster University Central Animal Facility. Mice were kept on a 12-hour light/12-hour dark cycle and provided access to water and food *ad libitum*.

### PCLS Generation and Culture

Animals were anesthetized with isoflurane and exsanguinated by severing the inferior vena cava. After sacrifice, the lungs were perfused by injecting 5mL of warm PBS into the right ventricle of the heart to flush out residual blood. The trachea was cannulated and 1.3mL of 40°C 1.5% low-melting point agarose (Invitrogen) dissolved in Hanks’ Balanced Salt Solution (HBSS) was slowly infiltrated into the lungs via the cannula, followed by a 0.2mL bolus of air to ensure agarose reached the lower airways. During lung inflation, mice were kept on a heating pad to maintain a warm temperature to prevent premature gelling of agarose. After inflation, mouse bodies were transferred onto ice and left to cool for 30 minutes to ensure complete gelling of agarose prior to excision of lungs. Lungs were then carefully excised, and lobes were separated. Each lobe was then affixed to a specimen holder, externally embedded in 2% agarose, and individually sliced (500µm thickness) in HBSS using a Compresstome VF-510-0Z vibrating microtome (Precisionary Instruments; speed setting: 1.5, oscillation setting: 9). PCLS cores (2mm or 4mm diameter) were obtained from full slices using a tissue puncher. **Figure 1** depicts a schematic of experimental overview for the study.

**Figure 1.**
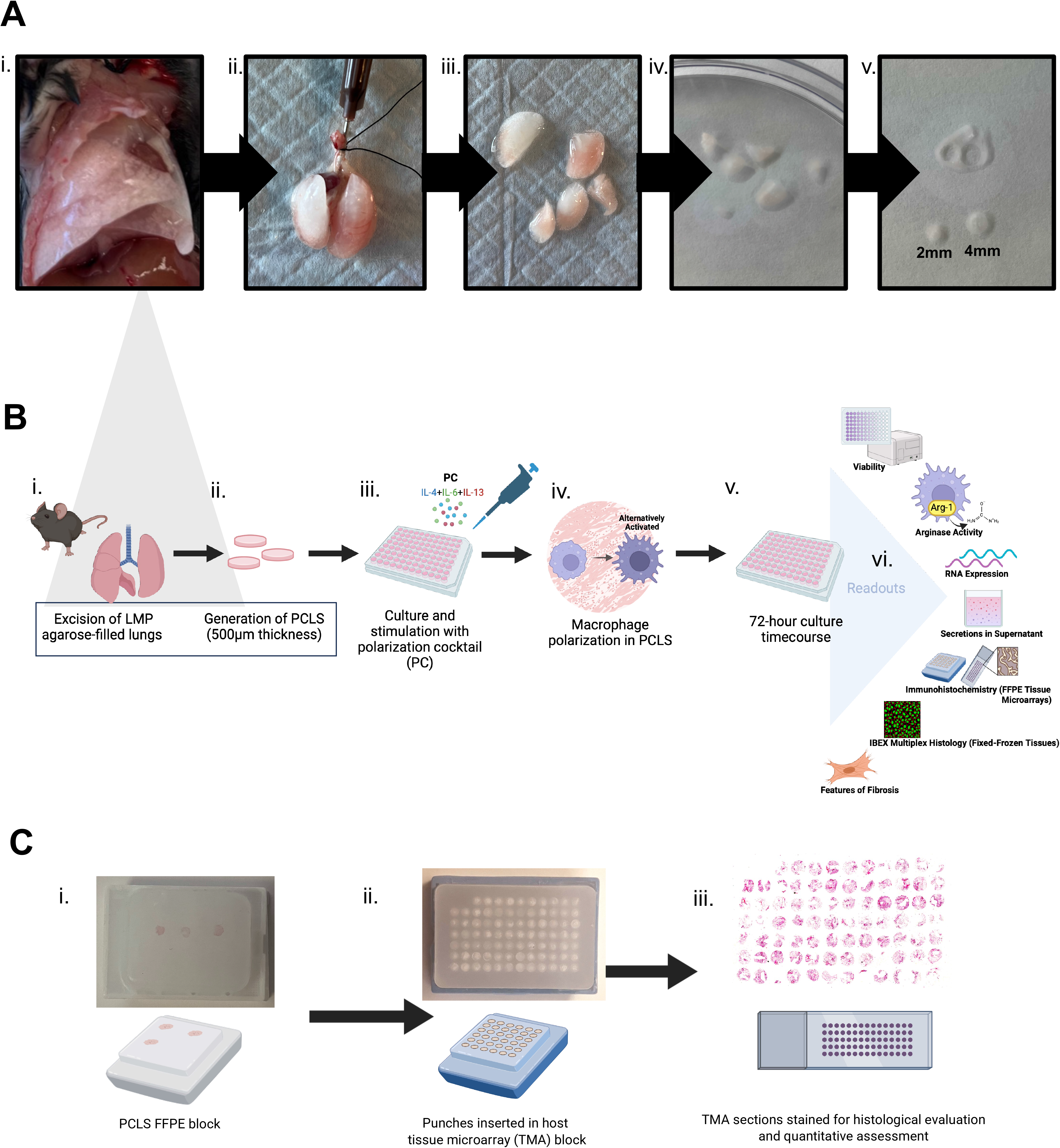
Schematic of overall experimental workflow. **(A)** Processing of murine lung for generation of PCLS. i. After sacrifice, lungs are infiltrated via the trachea with 1.5% low-melting point (LMP) agarose. Tissue is left to cool on ice in order for agarose to completely solidify prior to excision of lungs. ii. Filled lungs are excised from the body. iii. Lobes are separated to be sliced one at a time. iv. PCLS (500µm thickness) are generated from slicing each lobe using a Compresstome vibrating microtome. v. PCLS cores (2mm or 4mm in diameter) are punched from full lobe slices. **(B)** Experimental pipeline and readouts. i.,ii. Lungs are removed, sliced, and cored to create PCLS. After slicing, PCLS are placed in culture medium and left in incubator overnight to acclimate prior to treatment. iii. PCLS are moved to 96-well plates and treated with polarization cocktail (PC; IL-4+IL-6+IL-13). Baseline samples can also be harvested at this time (Day 0). iv.,v. Macrophage polarization occurs over 72-hour time course. Samples are harvested at 24-hour intervals throughout the time course. vi. Various readouts can be conducted on the polarized and control PCLS, including viability assays (WST-1), arginase activity assay, RNA expression, measurement of secreted components in supernatant, traditional brightfield immunohistochemistry (on FFPE PCLS, in tissue microarrays), multiplex immunostaining with Iterative Bleaching Extends Multiplexity method (IBEX; on fixed-frozen PCLS), and evaluation of fibrotic features (determined via histology, RNA expression, and secretions in supernatant). **(C)** Tissue microarray (TMA) generation from FFPE PCLS for histological evaluations. i. Original FFPE blocks containing PCLS. ii. 2mm punches are taken and inserted in parent TMA blocks. iii. Sections are taken from TMA and stained for histological evaluations, including quantitative assessment with HALO Image Analysis Software. Note: After fixation and prior to embedding, a small amount of eosin can be added to the ethanol storage solution to lightly colour the tissue, increasing ease of visibility during the embedding process. Figure created using BioRender.com.

After slicing, PCLS were cultivated in DMEM culture medium supplemented with 10% fetal bovine serum, 2 mM L-Glutamine, 100 U/mL penicillin, 100 µg/mL streptomycin, and 2.5 µg/mL amphotericin B at 37°C, 5% CO2. Medium was changed 3 times and PCLS were left in the incubator overnight to acclimate prior to treatment. At the time of treatment, PCLS were moved to 96-well plates and treated with a polarization cocktail (PC) consisting of recombinant IL-4 (40 ng/mL), IL-6 (10 ng/mL), and IL-13 (100 ng/mL; PeproTech), or a control cocktail (CC) consisting of only medium and diluent. We have previously shown that this triple cytokine combination stimulates profibrotic hyperpolarization of macrophages *in vitro* [24], however for *ex vivo* purposes we increased concentrations (while maintaining the same component ratios). Samples were harvested at 24-hour intervals over a 72-hour time course. For clodronate experiments, PCLS were subjected to a 24-hour pre-treatment period with liposomal clodronate (10-3000µM; Encapsula NanoSciences), followed by CC or PC treatment for 48 hours.

### Water-Soluble Tetrazolium-1 (WST-1) Assay

PCLS (4mm diameter) were incubated (37°C, 5% CO2) with 10µL of WST-1 reagent (Roche) in 100µL of culture media for 1 hour. After incubation, supernatant was transferred to a fresh 96- well plate and absorbance was measured at 450nm using a plate reader.

### Tissue Microarray Generation and Brightfield Histology

PCLS (2mm diameter) were fixed in 10% neutral-buffered formalin for 24 hours and then transferred to 70% ethanol. Fixed tissues were embedded in paraffin. Tissue microarrays were generated from the formalin-fixed paraffin embedded (FFPE) PCLS with the TMA Master II (3D Histech Ltd), by taking 2mm punches (containing full PCLS) from the original blocks and inserting them in a host paraffin block. FFPE histology was conducted at the John Mayberry Histology Facility at the McMaster Immunology Research Centre. 5µM sections were cut using a microtome (Leica) and stained with hematoxylin and eosin (H&E) and immunohistochemical (IHC) stains for Arginase-1, CD206, and α-SMA. Antibody information can be found in **Supplementary Table 1.** IHC was completed using a Bond RX immunostainer (Leica). Full slides were digitized (20X brightfield) using an Olympus VS120 Slide Scanner (Evident Scientific) at the Firestone Molecular Imaging and Phenotyping Core Facility (MPIC). Histological quantification was performed using HALO image analysis software (version 3.5, Indica Labs). For Arginase-1, CD206, and H&E, the Multiplex IHC module was used. For α-SMA, airways and vessels were excluded and the Area Quantification module was used.

### Iterative Bleaching Extends Multiplexity (IBEX) Fluorescent Histology

Multiplex staining of PCLS was achieved using IBEX, an open-source method for serial multi-marker immunolabelling [25,26]. PCLS (2mm diameter) were fixed in Cytofix/Cytoperm (BD Biosciences) diluted 1:4 in PBS, for 24 hours at 4°C. Tissue was then transferred to a 30% sucrose solution for cryoprotection for 48 hours at 4°C. Following fixation and cryoprotection, PCLS were embedded in optimal cutting temperature (OCT) compound and frozen using liquid nitrogen and isopentane. 12µM cryosections of fixed-frozen tissue were cut using a cryostat (Leica) and placed in chambered cover glasses. Tissues were incubated with the primary antibody staining solution overnight at 4°C. When needed, the secondary antibody staining solution was applied for 1 hour at 37°C. Antibody information can be found in **Supplementary Table 2.** After staining, Fluoromount-G mounting medium (Southern Biotech) was added. Multiplex imaging (20X fluorescent) was performed using an inverted confocal microscope (Zeiss LSM 980) at the McMaster University Centre for Advanced Light Microscopy (CALM). Tile images of the entire tissue were captured using a Plan-Apochromat 20X objective (0.8 numerical aperture), with pixel dimensions of 0.124um x 0.124um and a pinhole size of 1 Airy unit. Following image acquisition, mounting medium was removed and fluorophores were bleached by applying a 1mg/mL solution of lithium borohydride (Sigma-Aldrich) for 15 minutes. Tissues were stained with the next round of antibodies and the procedure was repeated iteratively as described. For histological analysis, serial image rounds were merged in HALO image analysis software, using DAPI as the alignment fiducial channel. For α-SMA staining, airways and vessels were excluded to focus on parenchymal expression. Quantification was performed using the Highplex FL module.

### Arginase Activity Assay

PCLS (4mm diameter) were homogenized in lysis buffer (0.1% Triton-X supplemented with sodium orthovanadate, PMSF, DTT and bovine lung aprotinin) using a Bullet Blender Bead Homogenizer (Next Advance). Homogenates were then centrifuged at maximum speed to remove remaining insoluble debris, and the arginase activity assay was carried out as previously described [7]. Briefly, lysates were diluted with 25mM Tris-HCl to form a 1:1 mixture, from which 25μL was transferred to a 96-well PCR plate containing 2.5μL 10 mM manganese chloride per sample well. The plate was then incubated in a thermal cycler at 56°C for 10 minutes. 25μL of 0.5M L-arginine then added, followed by another thermal cycler incubation at 37°C for 30 minutes. Urea standards, 200μL of sulfuric+phosphoric acid solution, and 10μL of 9% alpha- isonitrosopropiophenone were added to the plate. A final thermal cycler incubation was completed at 95°C for 30 minutes. After allowing the plate to cool for 5 minutes, 150μL from each well was transferred to a fresh 96-well flat bottom plate and absorbance was read at 550nm using a plate reader.

### Enzyme Linked Immunosorbent Assay (ELISA)

YM1 and CCL17 protein levels were measured in PCLS (4mm diameter) supernatant using commercially available ELISA kits (R&D Systems), according to the manufacturer’s protocol. Of note, CCL17 levels that were too low to be detected were assigned the lower limit of detection.

### RNA Isolation and cDNA Synthesis

PCLS (4mm diameter) were snap frozen using liquid nitrogen and were stored at −80°C until extraction. PCLS RNA isolation protocol was adapted from previously published studies [27,28]. Six PCLS per condition were pooled and homogenized in TRIzol Reagent (Invitrogen) using a bead mill. Phase separation was completed with chloroform and gel density tubes (Qiagen). The aqueous phase was then collected and RNA was extracted using the RNeasy Fibrous Tissue Mini Kit (Qiagen). RNA was converted to cDNA via reverse transcription using qScript cDNA SuperMix (Quantabio) according to the manufacturer’s instructions.

### Quantitative Real-Time Polymerase Chain Reaction (PCR)

PCR was performed using the QuantStudio 3 system (Applied Biosystems), with Advanced qPCR Mastermix (Wisent) and ThermoFisher Scientific Predesigned TaqMan Gene Expression Assay primer pairs (information found in **Supplementary Table 3**). Using *GAPDH* as the reference gene, relative expression was calculated as 2^-ι1CT^.

### Sircol Soluble Collagen Assay

The Sircol Soluble Collagen Assay 2.0 (Biocolor) was used to measure secreted soluble collagen in PCLS (4mm) supernatant, according to the manufacturer’s instructions.

### Statistical Analysis

Results were expressed as the mean ± standard error of the mean (SEM). Comparisons of two groups were performed with an unpaired two-tailed t test, while more than two groups were compared with ANOVA followed by Sidak’s multiple comparisons test. Statistical analyses were performed using GraphPad Prism 10. A P value less than 0.05 was considered statistically significant.

## RESULTS

### Development of Optimized PCLS Experimental Workflow

In the development of our experimental workflow, our objective was to establish a robust moderate-throughput approach to lung macrophages in an *ex vivo* environment. To maximize our throughput capacity, as well as normalize the size of PCLS, we created punches (2mm or 4mm in diameter) from full-lobe slices using a handheld tissue coring tool. We found punching the cores post-slicing, as opposed to taking cores from pre-sliced lobes, allowed for better control over the uniformity of core size. **Figure 1A** contains a photographic depiction of the workflow for processing murine lungs for PCLS generation. We have also included a summary of potential technical issues faced during the slicing process and troubleshooting solutions in **Supplementary Table 4.** On average, from slicing all lobes of a murine lung we are able to obtain approximately 40 4mm PCLS or 70 2mm PCLS (500uM thickness) per mouse (taking approximately 1.5 to 2 hours from sacrifice to incubator). In developing means to increase throughput of our PCLS studies, we introduced a tissue microarray-based approach for FFPE histology (**Figure 1C**). While previous studies have performed fundamental histological assessments on PCLS, these analyses have generally been limited in throughput. Constructing tissue microarrays from PCLS allows us to achieve this and evaluate approximately 80 PCLS cores (2mm diameter) on a single slide and using a tissue thickness of 500uM maximizes the quantity of FFPE serial sections that can be used for histological staining. Additionally, extracting adequate amounts and quality of RNA from PCLS has been reported as a challenge due to limitations of small tissue mass and interference from agarose [27,29]. Using a modified protocol derived from Stegmayr et al. (2020) [27] and Michalaki et al. (2022) [28], we isolated RNA using 6 pooled PCLS per condition (see Methods).

For the purpose of gene expression readouts via PCR, we obtained adequate RNA quantity and quality, from both control and PC-treated samples (**Supplementary** Figure 1).

### Murine Precision-Cut Lung Slices (PCLS) Remain Viable and Structurally Intact Throughout Culture Time course with Polarization Cocktail (PC) Treatment

First, to assess the utility and viability of our PCLS experimental system and treatment cocktail for subsequent macrophage polarization studies, we performed metabolic and histological evaluations at 24-hour intervals throughout the 72-hour culture period. Similar to previously published studies on human PCLS [22,30,31], we utilized the water-soluble tetrazolium (WST-1) assay and H&E histological staining to achieve this. Over the 72-hour time course in culture, PCLS maintained high viability and stable total cell numbers. Viability assessment of PCLS measured through mitochondrial activity using WST-1 assay (**Figure 2A**) demonstrated that CC- and PC-treated PCLS remained viable over 72 hours, and exhibited an increase in metabolic activity at the 48 and 72-hour timepoints, which may indicate tissue recovery after preparation [31]. Histological examination of H&E stained PCLS on an FFPE tissue microarray (**Figure 2B**), and quantification of cell number with HALO image analysis software (**Figure 2C**), confirmed that PCLS maintained structure and a consistent quantity of cells per total tissue area. No adverse effects from treatment with the PC were detected. Additionally, as our objective was to study and modulate macrophages in the PCLS system, we confirmed that macrophages persisted throughout the time course and were still present in the slices at the 72-hour timepoint, as also seen at baseline (0-hour) (**Figure 2D**). Overall, this allowed us to verify the integrity and suitability of our experimental system and PC intervention for investigation of *ex vivo* macrophage polarization.

**Figure 2.**
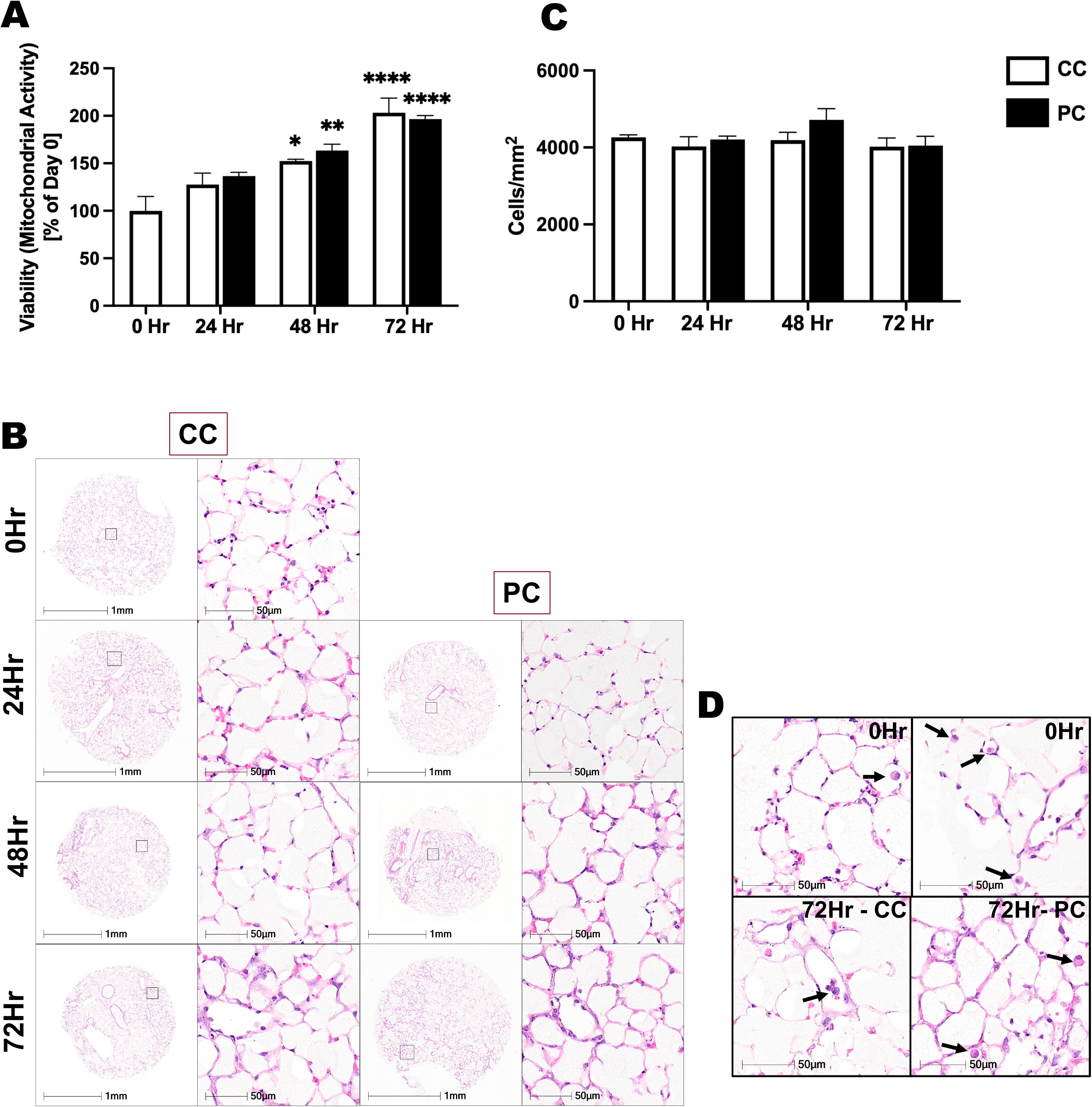
Murine Precision-Cut Lung Slices Maintain Viability and Structural Integrity in Culture Throughout Polarization Cocktail Treatment Time course. **(A)** Viability of precision- cut lung slices (PCLS) cultured with control cocktail (CC) or polarization cocktail (PC), assessed via water-soluble tetrazolium-1 (WST-1) assay. Absorbance values are expressed as percent of signal measured at Day 0. (n=3 mice, 5 slices per condition). **(B)** Hematoxylin and eosin (H&E) staining of PCLS (formalin-fixed, paraffin-embedded) throughout 72-hour time course. **(C)** Cell count per mm^2^ of tissue, quantified from whole slide images of H&E stained tissue microarrays using HALO image analysis platform (n=3 mice, 3-4 slices per condition). **(D)** Presence of macrophages in PCLS at baseline and 72-hour timepoint shown in H&E stained tissues. Arrows point to macrophages in alveolar spaces. * indicates P<0.05, ** indicates P<0.01, *** indicates P<0.001, and **** indicates P<0.0001, where * represents a significant difference compared to baseline (Day 0). Data are displayed as mean ± S.E.M.

### Treatment with the PC Induces Markers of Alternatively Activated Macrophages in PCLS Tissue and Supernatant

Next, we aimed to determine if the PC could polarize macrophages to an alternatively activated phenotype *ex vivo*, as we have previously shown in murine and human macrophages *in vitro* [24]. To investigate profibrotic macrophage polarization in the PCLS system, we began by measuring markers of profibrotic macrophages in the tissue (lysates) and soluble secretions (supernatant). Arginase-1, chitinase-3-like-protein-3 (YM1, gene name *Chil3*), CC chemokine ligand 17 (CC17), and cluster of differentiation 206 (CD206, gene name *MRC1*) are all established markers used to phenotype murine alternatively activated macrophages [32], which were found to be induced in PCLS with treatment of the PC (**Figure 3**). More specifically, arginase activity, determined by conversion of L-arginine to urea, was found to be increased in the lysates of PC-stimulated PCLS at all timepoints (24, 48, and 72 hours) during the time course, compared to CC-treated controls (**Figure 3A**). With regards to soluble markers, secreted YM1 and CC17 levels were increased with treatment of the PC at the 48 and 72-hour timepoints (**Figure 3B,C**), as measured by ELISA. At the gene expression level, treatment with PC led to increased gene expression of *Arg1*, *MRC1,* and *Chil3*, relative to *GAPDH*, at all timepoints throughout the time course (24, 48, and 72 hours; **Figure 3D-F**). Overall, these findings suggest the successful polarization of profibrotic pulmonary macrophages in the *ex vivo* PCLS environment, as exhibited through various quantitative assays.

**Figure 3.**
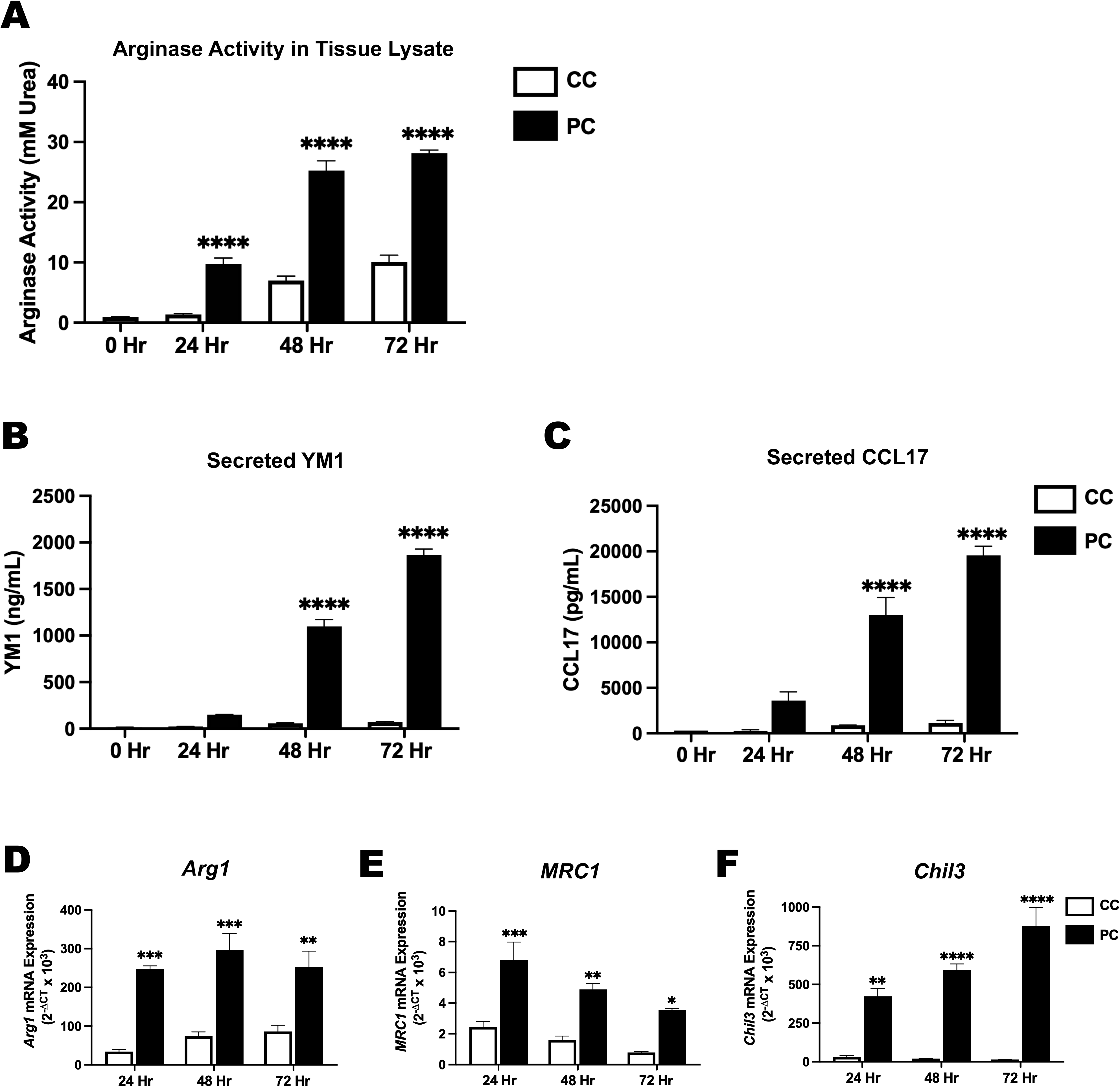
Treatment with the Polarization Cocktail Induces Markers of Alternatively Activated Macrophages in Precision-Cut Lung Slice Tissue and Supernatant. **(A)** Arginase activity measured as mM Urea in PCLS tissue homogenates (n=3 mice, 3-4 slices per condition). **(B,C)** Secreted Chitinase-3-like protein 3 (YM1) and CC chemokine ligand 17 (CCL17) protein levels in PCLS supernatant measured via ELISA (n=3 mice). **(D,E,F)** Normalized gene expression of Arginase 1 (*Arg1*), CD206 (*MRC1*) and YM1 (*Chil3*) in PCLS tissue, relative to *GAPDH* (n=3 mice, 6 slices pooled per condition). * indicates P<0.05; ** indicates P<0.01; *** indicates P<0.001; and **** indicates P<0.0001; where * represents a significant difference between the two treatment groups at the respective timepoint. Data are displayed as mean ± S.E.M.

### Histological Markers Characteristic of Profibrotic Macrophages are Increased Throughout Polarization Time course

Following the observations outlined in Figure 3, we proceeded to directly visualize polarized macrophages in the cocktail-treated PCLS tissue. We elevated the throughput capacity of our PCLS histological readouts through the construction of tissue microarrays containing numerous FFPE PCLS cores (see Methods), which could then be stained and analyzed on a single slide. To quantitatively assess alternatively activated macrophages throughout the time course, brightfield immunohistochemical (IHC) staining was performed on the tissue microarrays and whole-slide images were acquired. We observed that IHC staining for established alternatively activated macrophage markers, Arginase-1 and CD206, were increased with treatment of the PC (**Figure 4A,B**). Specifically, using HALO quantification we found that the percentage of positive cells, as well as staining intensity, assessed by HALO H-Score which accounts for marker staining strength and proportion, were increased at all time course intervals for both Arginase-1 (**Figure 4C**) and CD206 (**Figure 4D**). In terms of cellular localization, it was observed that Arginase-1 and CD206 positive macrophages exist in both the alveolar lumen and interstitium of PCLS (**Figure 4E**). Overall, these findings complement the results displayed in Figure 3 and visually confirm the presence of profibrotic macrophage polarization with PC treatment, as well as demonstrate the utility of a moderate-throughput histological approach for PCLS assessments.

**Figure 4.**
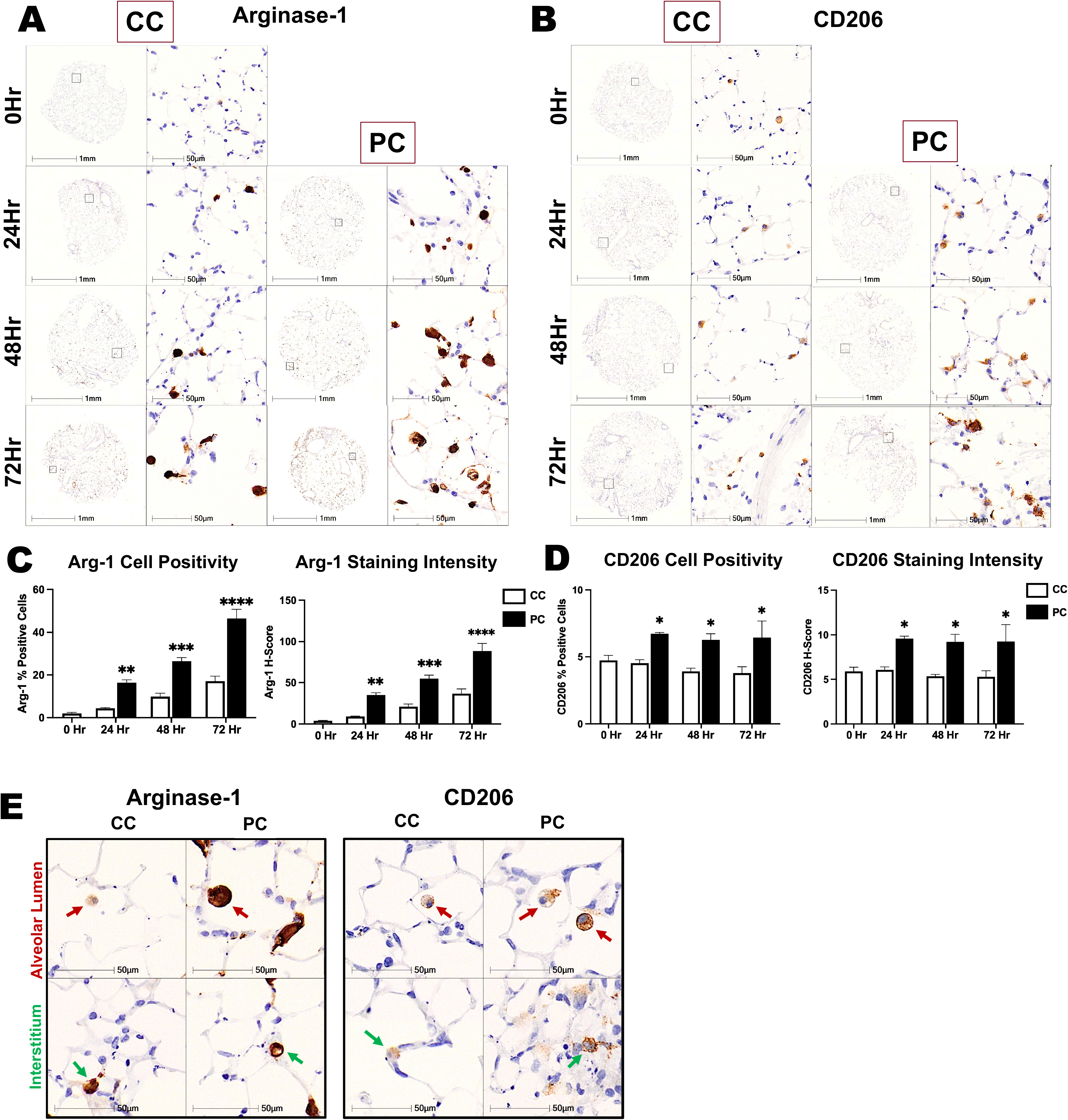
Histological Markers Characteristic of Profibrotic Macrophages are Increased Throughout Polarization Time course. **(A)** Representative images of Arginase-1 immunohistochemical (IHC) staining in CC- and PC-treated PCLS throughout time course. **(B)** Representative images of CD206 IHC staining in CC- and PC-treated PCLS throughout time course. **(C)** HALO quantification of Arginase-1 cell positivity and staining intensity (H-Score). **(D)** HALO quantification of CD206 cell positivity and staining intensity (H-Score). **(E)** Arginase- 1 and CD206 expression in macrophages located in both the alveolar lumen and interstitium, indicated by red and green arrows, respectively (72-hour timepoint). (n=3 mice, 3-4 slices per condition). * indicates P<0.05; *** indicates P<0.001; and **** indicates P<0.001; where * represents a significant difference between the two treatment groups at the respective timepoint. Data are displayed as mean ± S.E.M.

### PC Induces Polarization in Both Interstitial and Alveolar Macrophages, as Determined by Highly Multiplexed Staining (IBEX) to Assess Macrophage Phenotype

As the findings presented with traditional single-stain IHC established a foundation for characterizing the macrophages in the PC-stimulated PCLS, to better understand these cells we then sought to employ a high-content, multiplex imaging approach. Using Iterative Bleaching Extends Multiplexity (IBEX), an open-source method for serial multi-marker immunolabelling [25,26], we aimed to further investigate macrophage phenotype. Markers and antibodies used in our panel can be found in **Supplementary Table 2**. While Figure 4 conveys that Arginase-1 and CD206 IHC were increased in PCLS with treatment of the PC, and staining was present in cells that morphologically resemble macrophages, other cell types in the lung can also express these markers. To overcome the limitation of single-plex staining, we used IBEX to investigate macrophage-specific cells (defined as CD45^+^CD68^+^CD11c^+^) in the PCLS. First, to substantiate our findings, we examined overall profibrotic polarization (characterized by Arginase-1+CD206 expression) in the macrophage population (defined as CD45^+^CD68^+^CD11c^+^Arg1^+^CD206^+^ cells) (**Figure 5A**). Arg1^+^CD206^+^ macrophages were increased with PC treatment (48 hours), expressed as percentage of total cells in the PCLS (**Figure 5B**), as well as percentage of total macrophages (**Figure 5C**). Next, we aimed to evaluate if our PC could induce polarization in both interstitial (IM) and alveolar macrophages (AM), which are the two primary broad categories of macrophages found in the lung. In the context of lung fibrosis, both IM and AM are believed to play important roles in disease pathogenesis [33]. Delineating the exact contributions of IM and AM is a vital topic in the field under active investigation. Our brightfield IHC assessments, shown in Figure 4E, portray that Arginase-1 and CD206 positive cells were present in both the interstitium and alveolar lumen in PCLS. However, it is unclear if PC treatment definitively increases quantity of macrophage-specific polarization in each of these compartments. Thus, we aimed to evaluate polarization in IM and AM using IBEX. Murine IM and AM have been historically stratified based on the presence of CD11b and SiglecF expression, respectively [34]. We therefore used these markers, in combination with the general macrophage and polarization markers used in Figure 5A- C, to examine profibrotic polarization in IM and AM (**Figure 5D,E**). We observed that PC treatment (48-hours) increased the number of Arg1^+^CD206^+^ IM, expressed as percent of total IM (**Figure 5F**), as well as Arg1^+^CD206^+^ AM, expressed as percent of total AM (**Figure 5G**). Collectively, these results support the accumulated evidence that the PC successfully induces profibrotic macrophage polarization in our PCLS system.

**Figure 5.**
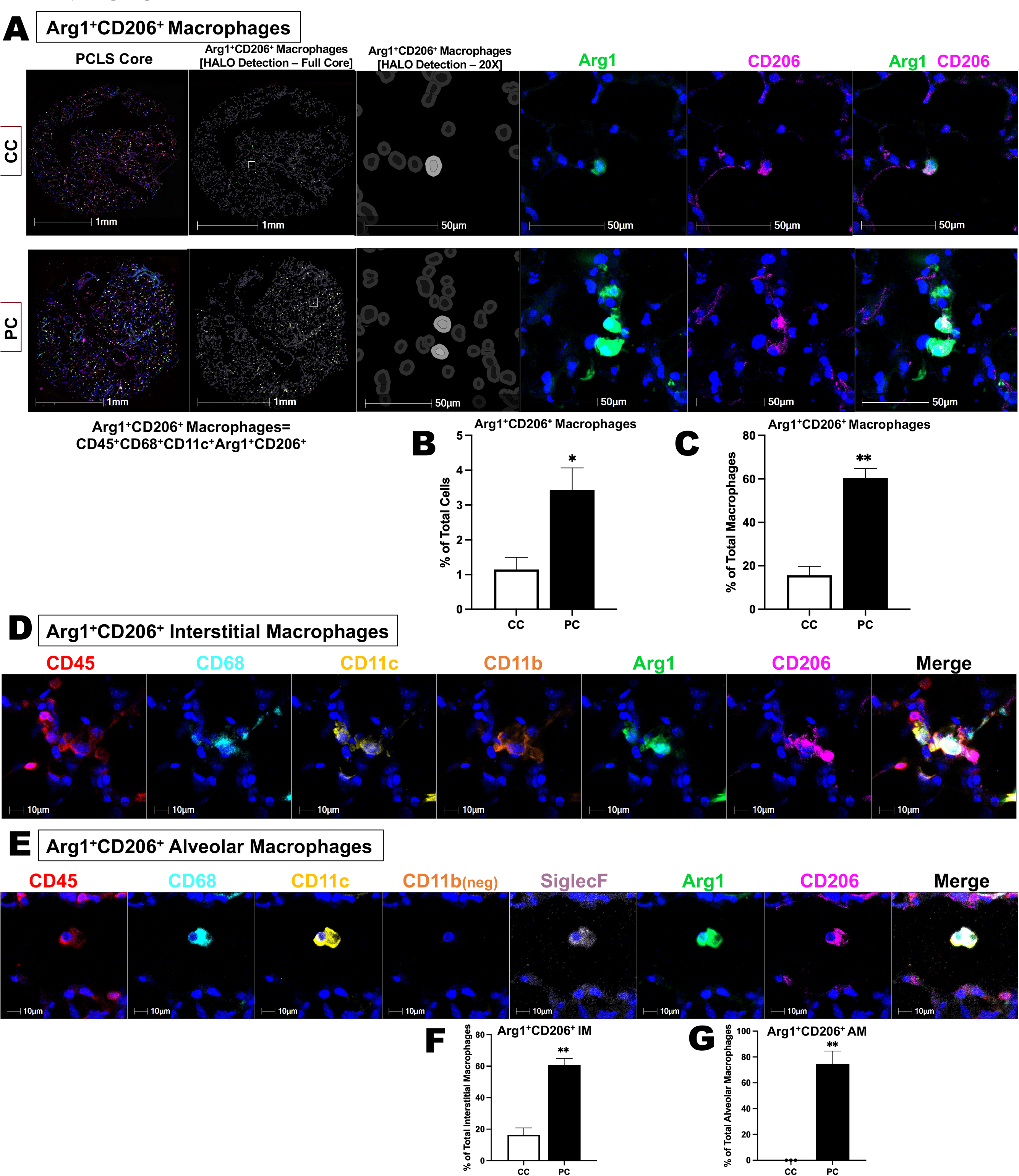
PC Induces Polarization in Both Interstitial and Alveolar Macrophages, as Determined by Highly Multiplexed Staining (IBEX) to Assess Macrophage Phenotype in PCLS. **(A)** Confocal fluorescent images of fixed-frozen multiplex stained (IBEX) PCLS treated with the CC and PC (48-hour timepoint). Markup for HALO detection of Arg1^+^CD206^+^ profibrotic macrophages (defined as CD45^+^CD68^+^CD11c^+^Arg1^+^CD206^+^ cells), as well as 20X representative images of Arginase-1 (green) and CD206 (magenta) staining in these cells, are shown. Cell nuclei are stained with DAPI (blue). **(B,C)** HALO quantification of Arg1^+^CD206^+^ profibrotic macrophages in CC- and PC-treated PCLS, expressed as percent of total cells and percent of total macrophages (n=3 mice). (**D,E)** Representative images of staining panels for Arg1^+^CD206^+^ interstitial macrophages (IM) (defined as CD45^+^CD68^+^CD11c^+^CD11b^+^Arg1^+^CD206^+^ cells) and Arg1^+^CD206^+^ alveolar macrophages (AM) (defined as CD45^+^CD68^+^CD11c^+^CD11b^-^ SiglecF^+^Arg1^+^CD206^+^ cells) in PCLS. Cell nuclei are stained with DAPI (blue). **(F)** HALO quantification Arg1^+^CD206^+^ IM, expressed as percent of total IM. **(G)** HALO quantification Arg1^+^CD206^+^ AM, expressed as percent of total AM. (n=3 mice). * indicates P<0.05; and ** indicates P<0.01; where * represents a significant difference between the two treatment groups. Data are displayed as mean ± S.E.M.

### Expression of Extracellular Matrix and Fibrotic Markers in PCLS Treated with PC

We have demonstrated the ability of the PC, consisting of IL-4, IL-13, and IL-6, to influence *ex vivo* macrophage polarization in PCLS. This is in alignment with *in vitro* studies that have shown the ability of these cytokines to hyperpolarize macrophages to the profibrotic phenotype [7,24]. Additionally, *in vivo* overexpression of IL-6 in the bleomycin lung fibrosis model induced an exacerbation of the fibrotic response, increased frequency of lung Arg1^+^CD206^+^ macrophages, and increased gene expression of IL-4 receptor in these macrophages (thus likely rendering them more susceptible to polarization by IL-4 and IL-13), demonstrating critical interplay between these cytokines, macrophages, and fibrosis [7]. While our primary objective in this study was to establish an *ex vivo* model suitable for studying pulmonary profibrotic macrophage polarization, given the key role of macrophages in fibrogenesis we speculated that it would be logical to investigate features related to fibrosis in our system. Using IHC staining of FFPE tissue microarrays, we evaluated the expression of α-SMA in PCLS subjected to CC or PC treatment (**Figure 6A**). HALO quantification of parenchymal α-SMA staining, excluding major airways and vessels, revealed an increase in α-SMA positive area at 48 and 72 hours in the PC-stimulated PCLS (**Figure 6B**). In the culture supernatant, we observed an increase in secreted soluble collagen with PC treatment at the 72-hour timepoint, as measured with Sircol Soluble Collagen Assay (**Figure 6C**). Normalized gene expression of α-SMA (*ACTA2*), extracellular matrix (ECM) component fibronectin (*FN1*), and ECM glycoprotein tenascin-C (*TNC*) were elevated in PC-treated PCLS lysates at the 72-hour (*TNC*), or both the 48 and 72-hour (*ACTA2* and *FN1*) timepoints (**Figure 6D-F**). Of note, gene expression of these three markers (*ACTA2, FN1, TNC*) has also been shown to be increased in a published model of fibrosis in human PCLS [22]. Lastly, to better understand the phenotype α- SMA^+^ cells in the PCLS, we performed multiplex image analysis with IBEX. In PC-treated PCLS, the α-SMA^+^ cell population in the parenchyma had increased co-expression of the marker profile of Arg1^+^CD206^+^ profibrotic macrophages (**Figure 6 G,H**). These results suggest that in addition to augmenting macrophage polarization, the PC may also have fibrosis-inducing properties in our experimental system.

**Figure 6.**
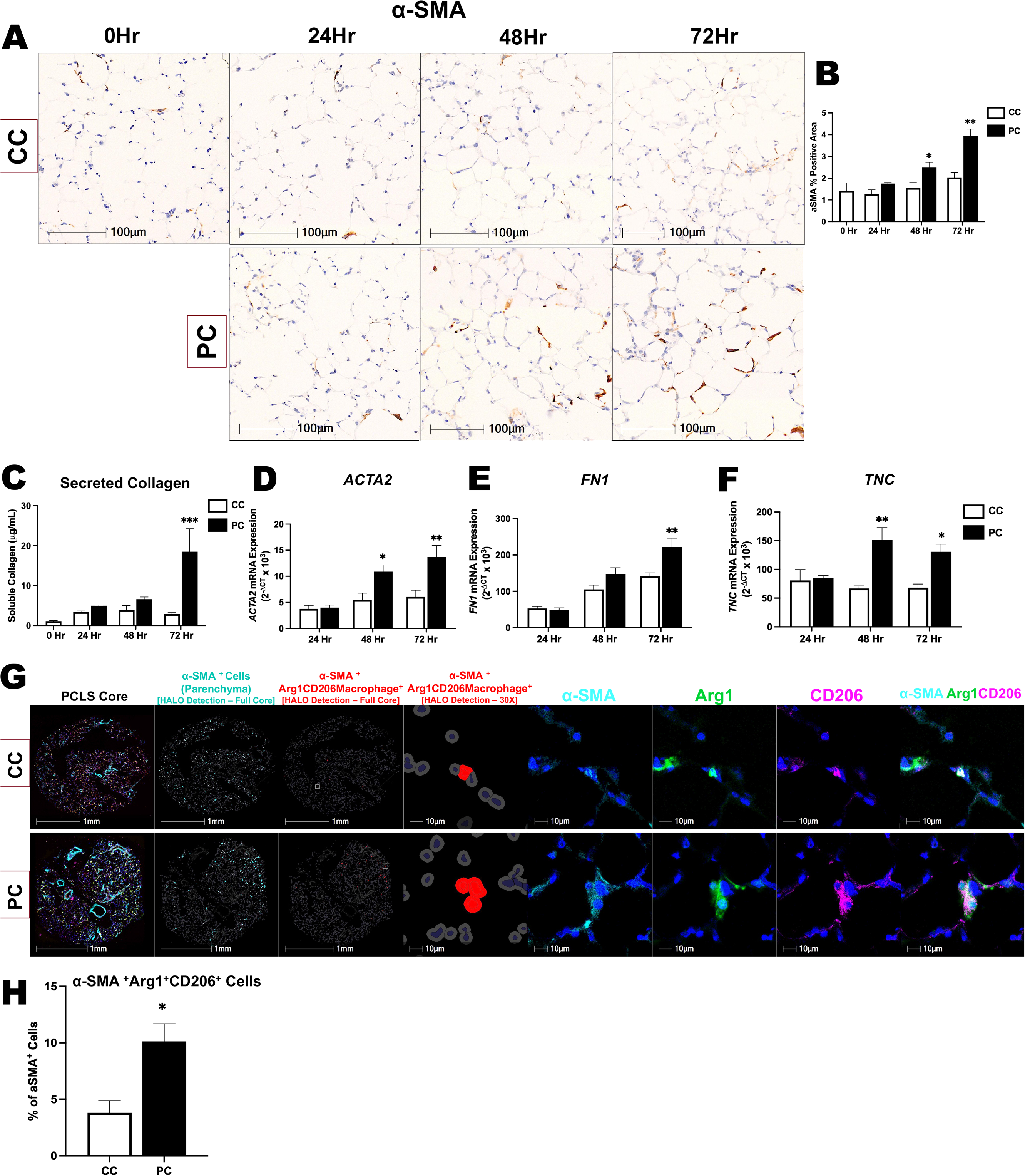
Expression of Extracellular Matrix and Fibrotic Markers in PCLS Treated with PC. **(A)** α-SMA IHC staining in CC- and PC-treated PCLS throughout time course. **(B)** HALO quantification of α-SMA positive parenchymal area (n=3 mice, 3-4 slices per condition). **(C)** Secreted soluble collagen in PCLS supernatant measured with Sircol Soluble Collagen Assay (n=3 mice). **(D,E,F)** Normalized gene expression of α-SMA (*ACTA2*), Fibronectin (*FN1*) and Tenascin- C (*TNC*) in PCLS tissue, relative to *GAPDH* (n=3 mice, 6 slices pooled per condition). **(G)** Confocal fluorescent images of multiplex stained (IBEX) PCLS treated with the CC and PC (48- hour timepoint). Markup for HALO detection of α-SMA+ cells (parenchyma), and cells that co- express α-SMA and markers for Arg1^+^CD206^+^ macrophages, is shown. 30X magnification images (CC- and PC-treated PCLS) of α-SMA (turquoise), Arg1 (green) staining, and CD206 (magenta) staining are also shown. Cell nuclei are stained with DAPI (blue). **(H)** HALO quantification of α- SMA and Arg1^+^CD206^+^ profibrotic macrophage marker co-expression in CC- and PC-treated PCLS, expressed as percent of α-SMA^+^ cells in the parenchyma (n=3 mice). * indicates P<0.05; ** indicates P<0.01; and *** indicates P<0.001; where * represents a significant difference between the two treatment groups at the respective timepoint. Data are displayed as mean ± S.E.M.

### Clodronate Treatment Diminishes Effects of PC on Profibrotic Macrophage Readouts

Finally, in order to substantiate the role of macrophages in the observed responses in the PCLS, we conducted a depletion study using liposomal clodronate. Clodronate is known for its ability to selectively deplete macrophages both *in vitro* and *in vivo* [35], and has been used in murine bleomycin studies to demonstrate that macrophages are required for lung fibrosis [36]. PCLS were pre-treated with liposomal clodronate for 24 hours, followed by 48 hours in culture with the CC or PC. Confirming clodronate-mediated depletion, macrophage quantity was decreased in PCLS, determined by counting visible alveolar macrophages in H&E stained tissue (**Figure 7A,B**). We next evaluated expression of profibrotic macrophage markers that are known to be induced by the PC. We observed that clodronate pre-treatment diminished PC-induced arginase activity in a dose-dependent manner (**Figure 7C**). Similarly, secreted YM1 (**Figure 7D**) and CCL17 (**Figure 7E**) in PCLS supernatant exhibited a dose-dependent decrease. In IHC-stained FFPE PCLS, clodronate dose-dependent reductions in Arginase-1 (**Figure 7F,G**) and CD206 (**Figure 7H,I**) positive cells and staining intensity (H-Score) were identified. With regards to features related to fibrosis, we observed a reduction in soluble collagen with all doses of clodronate (**Figure 7J**). Additionally, there was a trend of reduction in α-SMA expression in the FFPE PCLS tissue (**Figure 7K,L**). Overall, our results suggest that macrophages have a key role in the profibrotic phenotype observed in our PCLS system.

**Figure 7.**
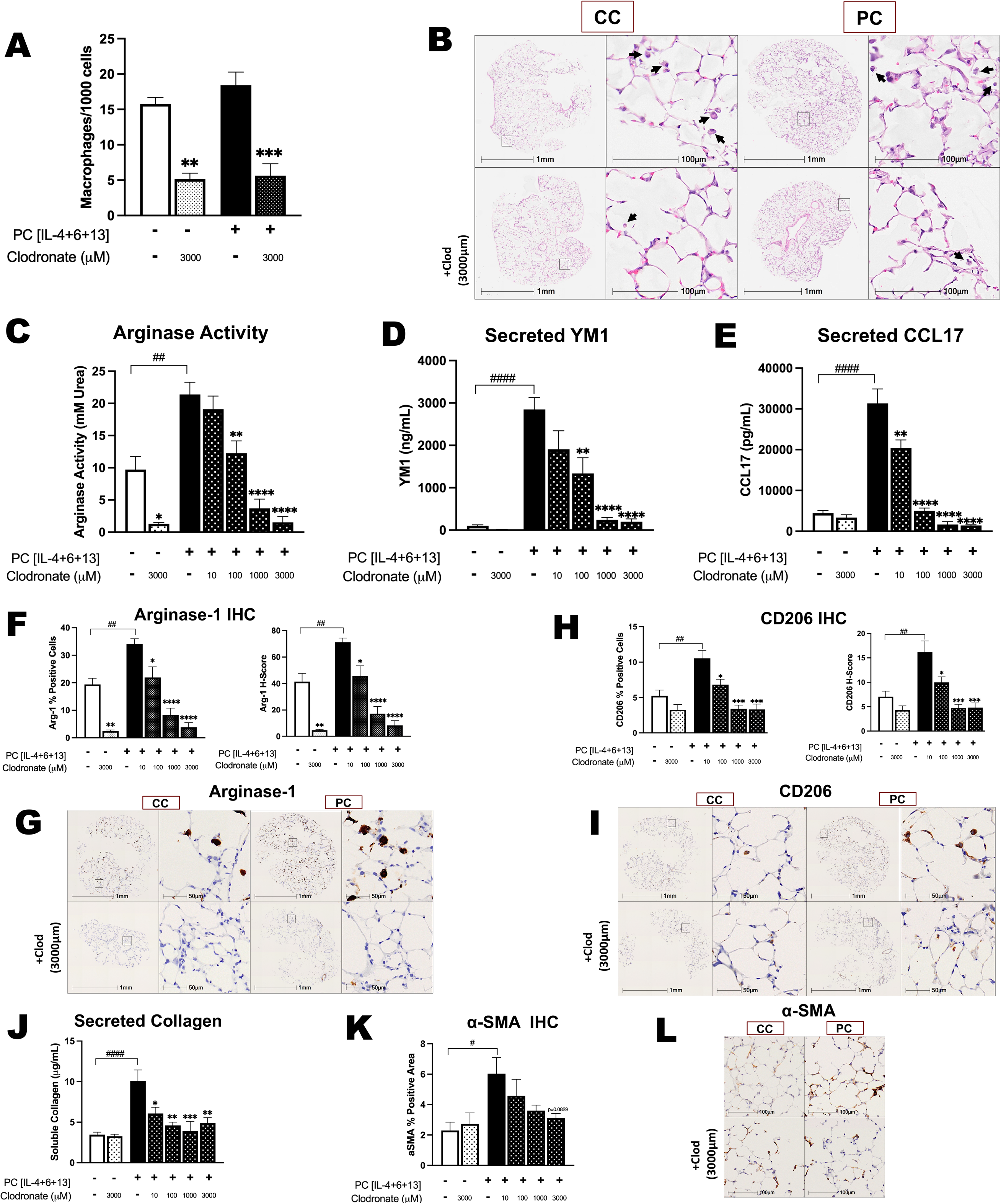
Clodronate Treatment Diminishes Effects of PC on Profibrotic Macrophage Readouts. PCLS were pre-treated with liposomal clodronate for 24 hours, followed by 48 hours in culture +/- PC. **(A)** Macrophage quantity per 1000 cells in PCLS, determined by counting visible alveolar macrophages in H&E stained tissue (n=3). **(B)** H&E images of CC- and PC-treated PCLS, +/- clodronate pre-treatment. Black arrows point to alveolar macrophages. **(C)** Arginase activity in PCLS homogenates treated with CC or PC, +/- clodronate pre-treatment (n=3 mice, 3 slices per condition). **(D,E)** Secreted YM1 and CCL17 protein levels in PCLS supernatant measured via ELISA (n=3 mice). **(F)** HALO quantification of Arginase-1 IHC cell positivity and staining intensity (H-Score). **(G)** Representative images of Arginase-1 IHC staining in CC- and PC-treated PCLS, +/- clodronate pre-treatment. **(H)** HALO quantification of CD206 IHC cell positivity and staining intensity (H-Score). **(I)** Representative images of CD206 IHC staining in CC- and PC-treated PCLS, +/- clodronate pre-treatment. (n=3 mice, 3-4 slices per condition). **(J)** Secreted soluble collagen in PCLS supernatant measured with Sircol Soluble Collagen Assay (n=3 mice). **(K)** HALO quantification of α-SMA positive parenchymal area (n=3 mice, 3-4 slices per condition). **(L)** Representative images of α-SMA IHC staining in CC- and PC-treated PCLS, +/- clodronate pre-treatment. *,# indicates P<0.05; **,## indicates P<0.01; *** indicates P<0.001; and ****,#### indicates P<0.0001; where * represents a significant difference between the clodronate pre-treated groups (patterned bars) and their respective CC or PC control group (solid-colored bars), and # represents a significant difference between the CC and PC groups (solid-colored bars). Data are displayed as mean ± S.E.M.

## DISCUSSION

Given the poor prognosis, debilitating symptoms, and gap in curative treatments for IPF, advancement of translational models and screening tools for preclinical therapies are especially critical. Macrophages are key regulators in tissue repair, however their plasticity and heterogeneity mean macrophage phenotypes are highly context-dependent, overall making the investigation of their precise contributions to disease challenging. In this study, we harness the potential benefits offered by the PCLS platform in establishing a novel strategy to investigate the profibrotic programming of lung macrophages, which have been demonstrated to be fundamental players in lung fibrosis [4–9]. In our model we have exhibited capacity for profibrotic macrophage programming with the treatment of a polarization cocktail, which is shown to be induced at multiple points throughout a 72-hour time course. This builds on the current literature revealing *ex vivo* immune competency in multiple PCLS disease models, including premalignancy, fibrosis, asthma, COPD, viral infection, bacterial infection, inflammation, and immunotoxicity [23,28,37–42]. Despite being disconnected from the circulating immune system, several studies have reported the presence and persistence of macrophages in PCLS. As demonstrated by Sompel et al. (2023), macrophages are the most abundant immune cell in murine PCLS [41]. Understandably, contribution to the lung macrophage population by incoming monocyte recruitment is absent in PCLS. Blomberg et al. (2023) have recently exhibited the capacity of macrophages to proliferate in a murine PCLS model of pre-cancer malignancy [21], which may contribute to the persistent presence of macrophages *ex vivo*. Additionally, scRNAseq analysis of cellular phenotype stability in PCLS demonstrated the preservation of immune cells including proliferating macrophages, alveolar macrophages, and monocyte-derived macrophages, although alveolar macrophage quantity was reported to be decreased after 120 hours of culture [20]. In the context of fibrosis, using scRNAseq analyses on human PCLS treated with the fibrosis cocktail developed by Alsafadi et al. (2017), Lang et al. (2023) showed induction of expression of fibrosis-related marker genes in a population of SPP1^+^ macrophages [22,23]. Taken together, our findings and the published evidence support the use of PCLS as a valid platform to investigate macrophage polarization and programming in the lung.

Previous findings from our group and others have shown that *in vitro* stimulation of human and murine macrophages with a combination of IL-4, IL-13 and IL-6 resulted in enhanced profibrotic programming to a hyperpolarized phenotype [7,24,43]. In the current study, we aimed to investigate if repurposing this cytokine combination (here termed the PC) could achieve similar polarization effects in an *ex vivo* setting. Similar to the published results, we observed an increase in profibrotic macrophage markers evoked with the PC in our *ex vivo* system, including Arginase- 1, CD206, YM1, CCL17, and arginase enzymatic activity [7,24,43]. To elicit profibrotic macrophage polarization, it is believed that the PC components work in synergy. IL-4 and IL-13 effectuate profibrotic macrophage polarization via IL-4 receptor alpha (IL-4Rα) signalling, and it has been shown that IL-6 upregulates IL-4Rα in macrophages *in vitro* [43,44], thus potentially increasing the propensity for polarization. It is plausible that similar processes are occurring with PC treatment *ex vivo*, however further investigations are required to delineate the molecular mechanism of action of profibrotic macrophage polarization in the PCLS.

Pulmonary macrophages are broadly categorized into IM and AM, which are both believed to participate in fibrogenic processes in lung fibrosis [9]. The delineation of their specific roles is a key topic of current investigation in the field, especially for IM which have been vastly understudied [33]. AM, residing in the lung alveoli, have been shown to alter tissue remodelling and interact with fibroblasts to enhance ECM synthesis [33,45]. IM, which are present in the lung interstitium, are less understood but are believed to be involved in initiation of fibrosis and partake in crosstalk with fibroblasts [33]. Circulating monocytes have been shown to contribute to the maintenance of both AM and IM, however both populations also demonstrate capacity for self-renewal [33]. In our PCLS model, we show the that the PC induces polarization in both IM and AM, thus demonstrating potential to be used as a tool to study these cells and their respective properties further. To target the specific role of IM in the relative absence of AM in future studies, AM could be largely washed out with repeated bronchoalveolar lavage fluid collection, prior to PCLS generation. Additionally, previous studies have shown retained functionality of these cells in PCLS, where IM were substantial antigen-uptaking cells of house dust mite extract *ex vivo* [46]. The primary objective for our study was to establish an *ex vivo* model for profibrotic macrophage programming in PCLS, using the PC. In addition to macrophage polarization, we interestingly observed other features related to fibrosis in our model, including ECM and α-SMA expression, particularly at later timepoints in the time course. The current paradigm of pathogenesis for pulmonary fibrosis is multifactorial and is believed to be initiated with insult to the lung epithelium. The resulting inflammatory response involves recruitment of macrophages to the site of injury, which through a series of mediators eventually cascades into activation of fibroblasts and resulting ECM deposition, resulting in fibrosis [9]. One plausible explanation for the observed fibrotic characteristics in our model is that the slicing of the lungs to generate PCLS functions as an injury, which then activates an intrinsic wound healing response in the tissue. This has been suggested previously [47], and may also explain why we see increased expression of some markers in our control PCLS, including *FN1*, Arginase activity, and Arg-1 IHC, compared to baseline. Providing external cytokines through administration of the PC further exacerbates the macrophage response and involvement, thus resulting in the development of fibrosis-like features in the PCLS.

Additionally, although shown to effect macrophages, the addition of the PC to PCLS culture does not solely target one cell type, and can influence a variety of cells. IL-4 and IL-13 have historically been shown to stimulate collagen synthesis in fibroblasts [48,49]. IL-4Rα signalling also plays a role in tissue remodelling, and has also been demonstrated to be essential for the development of lung fibrosis *in vivo* [50]. Thus, another potential explanation is direct modulation of fibroblasts by the PC, in addition to macrophages, cultivating fibrotic features. Furthermore, our group has previously shown that overexpression of IL-6 in a bleomycin-induced lung fibrosis model led to increased accumulation of profibrotic macrophages (Arg1^+^CD206^+^), as well as IL-4Rα expression, and worsened the fibrotic response [7]. This further supports the multifaceted interplay between the cytokines in the PC, polarized macrophages, and the fibrotic response. Lastly, the origin of α- SMA^+^ cells in fibrotic disorders remains a topic of debate [51]. Recently, there has been increasing evidence for macrophage to myofibroblast transition (MMT) in fibrosis, both in the lung and other organs, where α-SMA^+^ cells arise from the macrophage lineage and co-express markers for both cell types [51,52]. MMT processes may describe the increase in percentage of α-SMA^+^ cells co-expressing markers of Arg1^+^CD206^+^ macrophages in our system, and serve as a potential source of α-SMA^+^ cells.

Compared to traditional *in vitro* models, PCLS forgo the need for researcher-made recapitulation of the lung microenvironment, and present advantages of increased complexity, preserved architecture, and resident cell milieu, which substantiate their translatability in the study of lung pathology [53]. The *ex vivo* system is also less demanding of resources and time relative to *in vivo* models of disease, supporting utility especially in early stages of compound testing. Our goal was to establish a translational platform to study macrophage behaviour in fibrotic systems, with moderate-throughput that could be used for a variety of readouts. We introduce an approach to expand the throughput of direct histological evaluation in organ slices by constructing tissue microarrays, which can house approximately 80 PCLS that can be stained, imaged, and analyzed on a single slide. In the era of exponentially growing interest in spatial biology, there is also reasonable potential capacity for these analyses in cultured organ slice systems, which can be performed with the use of high-content imaging pipelines such as IBEX and other established methodologies. Future assessments using comprehensive gene profiling, such as RNAseq or microarray analyses, are also achievable with the attained RNA quantity and quality from our system. Additionally, multiple PCLS studies, including in the fibrosis field, have utilized human tissue to model disease and investigate disease mechanisms. These studies are highly robust and representative of human biology, as they are derived directly from human tissue. However, working with fresh human tissue presents logistical limitations, including inconsistencies in agarose filling, variability in baseline tissue viability, amount of tissue available, and/or lack of access to human tissue altogether. Murine models are therefore convenient tools to bypass these limitations, and can be used in synergy with human models to conduct screening on preclinical mechanisms and experimental treatment candidates. Overall, given the current set-up of our system, we believe our approach has potential for future expansion in several domains, including bleomycin pulmonary fibrosis models, PCLS generation from other species including humans, evaluation of pulmonary stretch and mechanotransduction, and testing of experimental treatments. The pipeline is scalable, and throughput could be increased further with the addition of multiple tissue slicers to simultaneously generate slices from multiple lobes, which is currently our time-limiting step.

In summary, we have established and validated a moderate-throughput, complex, viable model to study profibrotic polarization of macrophages in the lung using PCLS. Stimulation of PCLS with the PC effectively induces polarization of macrophages to a profibrotic phenotype. Our model provides means for interrogation of programming of profibrotic lung macrophages with the benefits of increased translatability, with future capacity for investigating novel therapeutic candidates and modes of action in pulmonary fibrosis.

## AUTHOR CONTRIBUTIONS

**MV:** methodology, formal analysis, investigation, writing – original draft, writing – review & editing, visualization; **AA**: methodology, investigation, writing – review & editing; **PA:** methodology, investigation, writing – review & editing; **VK:** methodology, writing – review & editing; **SN:** methodology, writing – review & editing; **TI:** methodology, writing – review & editing; **JK:** methodology, resources, writing – review & editing**; MK:** project administration, resources, supervision, writing – review & editing, funding acquisition; **KA:** conceptualization, resources, supervision, writing – review & editing, project administration, funding acquisition

## ACKNOWLEDGEMENTS

We would like to thank Mary Jo Smith, Mary Bruni, and Xiaoxing Ma at the McMaster Immunology Research Centre John Mayberry Core Histology Facility for their technical support in FFPE immunohistochemistry. We also thank Dr. Joao Pedro Bronze de Firmino and Dr. Mouhanad Babi from the McMaster Centre for Advanced Light Microscopy (CALM) for their expertise in confocal microscopy. We thank Vitoria Murakami Olyntho for knowledgeable teaching of the IBEX method. Lastly, we express our sincere thanks to Joanna Kasinska and Fuqin Duan for their expert technical laboratory assistance.

