## Supplementary Table 1, Supplementary Table 2, Supplementary Table 3, Supplementary Table 4, Supplementary Figure 1 for "A Novel *Ex Vivo* Approach for Investigating Profibrotic Macrophage Polarization Using Murine Precision-Cut Lung Slices"

**SUPPLEMENTARY MATERIAL**

**Supplementary Table 1: Antibody Information for FFPE IHC**

| Target Name | Clone | Host Species | Supplier | Catalog Number |
|---|---|---|---|---|
| α-SMA | 1A4 | Mouse | Agilent Dako | M0851 |
| Arginase-1 | D4E3M | Rabbit | Cell Signaling Technology | Cs93668 |
| CD206 | - | Rabbit | Abcam | ab64693 |

**Supplementary Table 2: Antibody Information for IBEX**

| Target Name | Clone | Host Species | Supplier | Catalog Number | Fluorophore |
|---|---|---|---|---|---|
| α-SMA | 1A4 | Mouse | Invitrogen | M0851 | eFluor660 |
| Arginase-1 | D4E3M | Rabbit | Cell Signaling Technology | Cs93668 | - |
| CD11b | M1/70 | Rat | BD Biosciences | 557397 | PE |
| CD11c | N418 | Hamster | Invitrogen | MCD11C20 | AF488 |
| CD206 | C068C2 | Rat | BioLegend | 141709 | AF488 |
| CD45 | 30-F11 | Rat | BioLegend | 103144 | AF594 |
| CD68 | FA-11 | Rat | BioLegend | 137004 | AF647 |
| Siglec F | E50-2440 | Rat | BD Biosciences | 552126 | PE |
| Anti-Rabbit | - | Goat | Invitrogen | A11008 | AF488 |

**Supplementary Table 3: Taqman PCR Primer Information**

| Gene | Catalog # (ThermoFisher Scientific) |
|---|---|
| *ACTA2* | Mm01546133_m1 |
| *Arg1* | Mm00475988_m1 |
| *Chil3* | Mm00657889_mH |
| *FN1* | Mm01256744_m1 |
| *GAPDH* | Mm99999915_g1 |
| *MRC1* | Mm00485148_m1 |
| *Tnc* | Mm00495662_m1 |

**Supplementary Table 4: Troubleshooting Issues in Generation of Murine PCLS**

| Issue | Potential Reasons | Solutions |
|---|---|---|
| Uneven filling of lung lobes | • Premature gelling of agarose in proximal airways | • Ensure LMP agarose is at a temperature of 40°C (no less than 37°C) directly prior to infiltration[1]<br>• Utilize heating pad and/or heat lamp to keep mouse body warm[1]<br>• Pour warm (37°C) HBSS on lungs directly prior to infiltration[1]<br>• Inject bolus of air (~0.2mL) into lungs to push agarose into distal airways[1] |
| Lungs are not fully inflated after agarose infiltration | • Premature gelling of agarose in proximal airways<br><br>• Too small of volume of agarose used for infiltration | • See above[1]<br><br>• Inject volume equivalent to total lung capacity of mouse (~1mL-1.3mL, depending on size of animal)[2] |
| Agarose leakage while filling lung | • Premature gelling of agarose in proximal airways, leading to blockage and subsequent rupture/damage<br><br>• Infiltration volume exceeded total lung capacity | • See above[1]<br><br>• See above[2] |
| Agarose leakage after filling lung | • Canula and/or syringe removed before agarose fully solidified<br><br>• Canula loose and/or ligature not secured tightly enough | • Leave canula and syringe in place while mouse body is on ice and agarose fully solidifies (30 minutes)<br><br>• Ensure depth of canula insertion in the trachea is not too shallow. Secure tightly with double-knotted ligature |
| Tissue is not being sliced by vibratome | • Inadequate infiltration of lungs with agarose<br><br>• Tissue is not properly secured to vibratome specimen holder | • See above[1,2]<br><br>• Ensure bottom side of tissue is completely glued to specimen holder. Do not allow glue to touch anywhere else on tissue |
| Tissue tearing while slicing | • Lung tissue is overfilled with agarose<br><br>• Inappropriate vibratome speed and/or oscillation | • See above[2]<br><br>• Lower vibratome speed and/or raise oscillation |

**Supplementary Figure 1: RNA Extraction Quantity and Quality Measures from Murine PCLS**

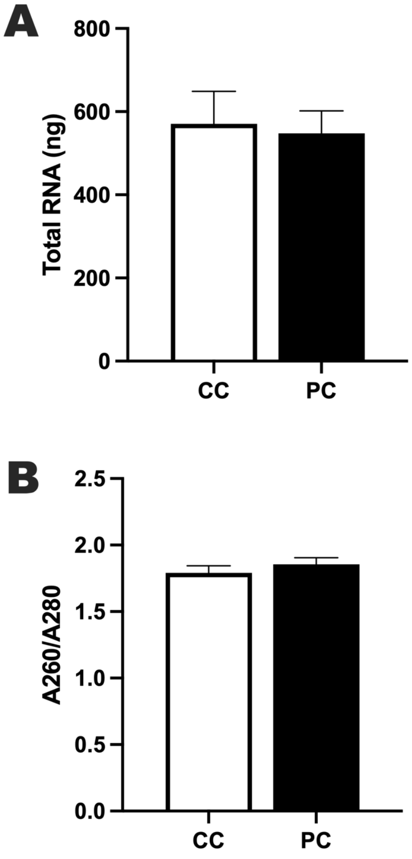


**Figure S1. RNA Extraction Quantity and Quality Measures from Murine PCLS.** Six PCLS (4mm diameter) were pooled for each sample. **(A)** Total RNA extracted from each sample. **(B)** A260/A260 ratio. n=18 PCLS from 3 mice. Results represent mean ± S.E.M. with samples taken from all timepoints.
